# Characterizing the response of *Acinetobacter baumannii* ATCC 17978 to azithromycin in multiple *in vitro* growth conditions

**DOI:** 10.1101/2020.05.19.079962

**Authors:** Nicholas Dillon, Hannah Tsunemoto, Saugat Poudel, Michael J. Meehan, Yara Seif, Richard Szubin, Connor A. Olson, Akanksha Rajput, Geovanni Alarcon, Anne Lamsa, Alison Vrbanac, Joseph Sugie, Samira Dahesh, Jonathan M. Monk, Pieter C. Dorrestein, Rob Knight, Adam M. Feist, Joe Pogliano, Bernhard O. Palsson, Victor Nizet

## Abstract

Multi-drug resistant (MDR) *Acinetobacter baumannii* is one of the most concerning pathogens in hospital infections. *A. baumannii* is categorized as an “Urgent Threat” by the U.S. Centers for Disease Control and the highest priority pathogen by the World Health Organization due to its propensity for broad antibiotic resistance and its associated high mortality rates. New treatment options are urgently needed for MDR *A. baumannii* infections. Our prior studies have demonstrated an unappreciated utility of the macrolide azithromycin (AZM) against MDR *A. baumannii* in tissue-culture medium. This finding is all the more surprising since AZM has no appreciable activity against *A. baumannii* in standard bacteriological media. The basis for this media-dependent activity of AZM against *A. baumannii* is not fully defined. In this study, we utilize a variety of techniques (growth dynamics, bacterial cytological profiling, RNA sequencing, and LC/MS) to profile the response of MDR *A. baumannii* to AZM in both standard bacteriological and more physiological relevant mammalian tissue-culture medium.

|  |  |
| --- | --- |
| Design type(s) | • replicate design • transcription profiling design • sequence analysis objective |
| Measurement(s) | • minimal inhibitory concentration • cDNA • transcription profiling assay • time-course exo metabolome • cellular morphology • bacterial growth • Genome sequence |
| Technology Type(s) | • RNA sequencing • fluorescence microscopy • liquid chromatography-tandem mass spectrometry • optical density measurement • Nanopore MinION • Illumina Hiseq 4000 |
| Factor Type(s) | • culture medium • biological replicate • experimental condition |
| Sample Characteristic(s) | • *Acinetobacter baumannii* • culturing environment |

## Background and Summary

*Acinetobacter baumannii* is an emerging human pathogen with a high propensity for broad antibiotic resistance. Initially gaining national attention due to its prevalence in US military personnel returning from the middle east, *A. baumannii* infections have now been detected worldwide^1^. The rapidity for which resistance has developed in *A. baumannii* especially considering it has only recently emerged as a predominate cause of ICU-acquired pneumonia, highlights the potential threat posed by this pathogen. Accordingly, *A. baumannii* has recently been upgraded to an “Urgent Threat” pathogen by the US Centers for Disease Control (CDC)^2,3^ and it is the #1 priority pathogen (critical priority) in the World Health Organization’s (WHO) “Global Priority Pathogens List for Combatting Antibiotic Resistance”^4,5^.

*A. baumannii* is a Gram-negative coccobacillus that is commonly classified as an opportunistic pathogen. Infections with *A. baumannii* can be acquired from both community and healthcare settings in addition to environmental sources^2^. *A. baumannii* infections are often detected in immuno-compromised patients, and present as either bloodstream or wound infections, or as the cause of bacterial pneumonia^6,7^. *A. baumannii* is understudied compared to other common MDR pathogens, and as such its virulence mechanisms are not well defined.

Approximately two-thirds of all *A. baumannii* infections are multi-drug resistant in the United States^3^. Resistance to carbapenems, mediated through the production of carbapenemases or metallo-β - lactamases, is prevalent leading to carbapenems no longer being a reliable treatment option for *A. baumannii*^2^. Owing to both target site mutations in the genes encoding DNA gyrase and topoisomerase IV, as well as enhanced efflux mechanisms, flouroquinolone resistance is also high in clinical *A. baumannii* isolates^2^. *A. baumannii* employs all three of the major mechanisms of horizontal gene transfer (conjugation, transformation, and transduction) to aid in its acquisition and spread of various antibiotic resistance mechanisms^8^.

High rates of resistance^2^, combined with a failing drug delivery pipeline^9^, necessitate a push to identify new efficacious therapeutical options for MDR *A. baumannii* infections. While pursuing potential new therapies we uncovered unappreciated activity for the macrolide azithromycin (AZM) against MDR *A. baumannii*^10,11^. Interestingly, enhanced AZM activity was found when *A. baumannii* was cultured in amended Roswell-Park memorial institute (RPMI+) tissue culture medium but not in the standard bacteriologic medium cation-adjusted Mueller-Hinton broth (CA-MHB)^10,11^. The basis for the media dependent differences in AZM susceptibility have not been fully defined.

In this study we are exploring the basis for the media dependent changes in AZM susceptibility for *A. baumannii*. For this analysis we have selected *A. baumannii* strain ATCC 17978 from the American type culture collection. Strain ATCC 17978 was originally isolated from a fatal meningitis in a pediatric patient, and is an established drug-sensitive strain of *A. baumannii*^12^. Strain ATCC 17978 was cultured in either CA-MHB or RPMI+ and exposed to varying concentrations of AZM. Growth was monitored and samples were collected for analysis via bacterial cytological profiling^13^, DNA sequencing and genome assembly, RNA sequencing, and untargeted liquid chromatography mass spectrometry data acquisition.

## Methods

The methodology and approach used in this study were based our previously published articles with *Staphylococcus aureus*^14–16^ but adapted for *A. baumannii*.

### Bacterial Strain and Culturing Reagents

*Acinetobacter baumannii* strain 17978 was acquired from the American type culture collection (ATCC) and is herein denoted strain ATCC 17978. Mueller-Hinton broth (MHB) (Sigma-Aldrich) was supplemented with 25 mg/L Ca^2+^ and 12.5 mg/L Mg2+ (CA-MHB) was used as the standard bacteriological medium throughout the experiments. The tissue culture media Roswell-Park memorial institute medium 1640 (RPMI) (Thermo Fisher Scientific) supplemented with 10% Luria broth (LB) (Criterion) (RPMI+) was used in the experiments to represent a more physiologically relevant bacterial culture medium.

### Antibiotic Susceptibility Testing and Growth Conditions

Minimum inhibitory concentration (MIC) experiments were carried out as previously described^11^. Briefly, AZM (Fresenius Kabi) was serially diluted in the associated medium to generate working stocks of the drug. *A. baumannii* ATCC 17978 cultures were taken out of −80 °C storage and fresh cultures were inoculated and grown overnight. New cultures were started from the overnight cultures the following morning, grown until mid-log phase at an OD_600_=~0.4 (~1*10^8^ CFU/mL). The mid-log phase ATCC 17978 culture was then used to inoculate the experimental cultures (+/- AZM) at a starting OD_600_=~0.01 (~2.5*10^7^ CFU/mL). These experiments utilized 3 biological replicates and growth was monitored every 30 min for 3 hours. Growth in the presence of AZM in either CA-MHB or RPMI+ was recorded and plotted (Fig. 1).

**Figure 1.**
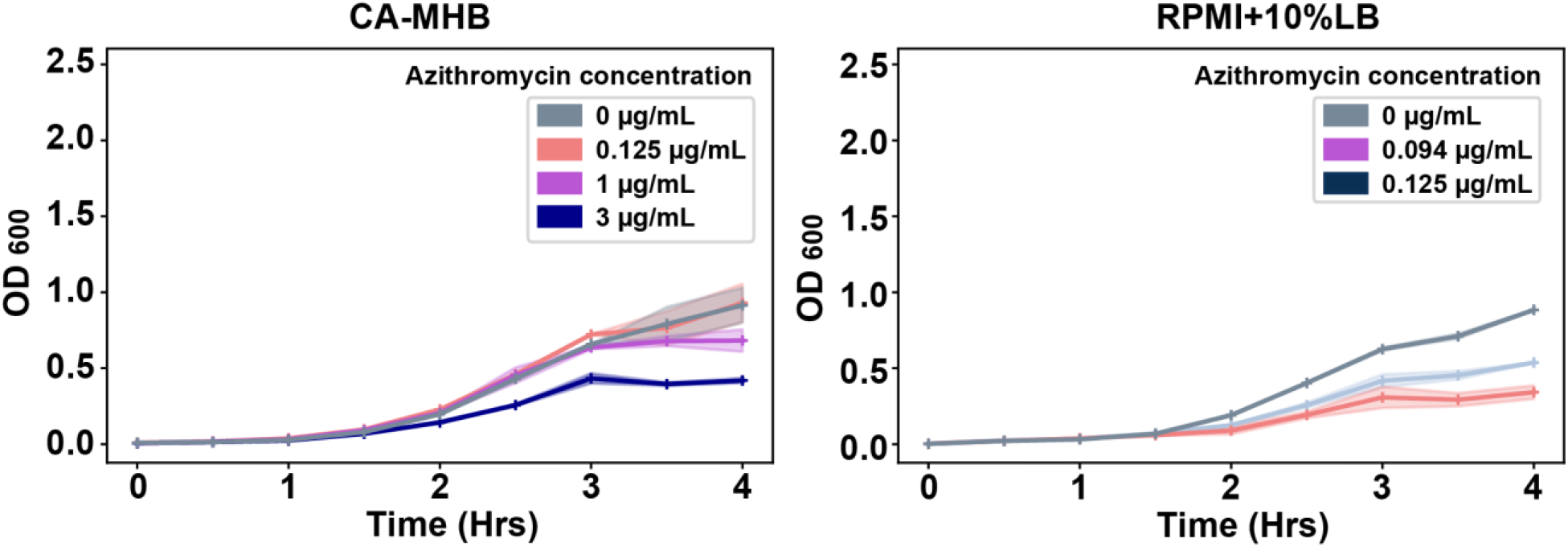
*Acinetobacter baumannii* strain ATCC 17978 growth in the presence of azithromycin at various sub-inhibitory concentrations in CA-MHB and RPMI + 10% LB media.

### Bacterial Cytological Profiling

Fluorescence microscopy was done as previously described with modifications^11,14,15,17^. In brief, at the 2.5 h timepoint, 20 μL cells were added to 1 μL dye mix containing 80 μg/mL DAPI, 10 μM SYTOX Green, and 200 μg/mL FM4-64 in 1x T-base. 6 uL sample was then transferred to an agarose pad (20% media, 1.2% agarose) on a single-well glass slide and imaged using an Applied Precision DV Elite epifluorescence microscope with a CMOS camera. The exposure times for each wavelength were kept constant for all images and were as follows, TRITC/Cy-5 = 0.05s, FITC/FITC = 0.01s, DAPI/DAPI = 0.05s.

Deconvolved images were adjusted using both FIJI (ImageJ 1.51w) and Adobe Photoshop (2015.1) to remove the background in FM4-64 and DAPI channels. This was done to ensure that cell and DNA objects are within the highest intensity quartile. Adjusted images were then run through a custom CellProfiler 3.0 pipeline that individually thresholds and filters FM4-64 and DAPI channels to obtain segmentation masks for the cytological features cell wall, DNA, and entire cell. Identified objects were further processed in CellProfiler to obtain a total of 5,285 features^18,19^. Feature selection was necessary to create a subset of physiologically relevant features to minimize redundancy within the dataset prior to analysis. Fig. 2 presents a summary of processing steps required for the bacterial cytological profiling.

**Figure 2.**
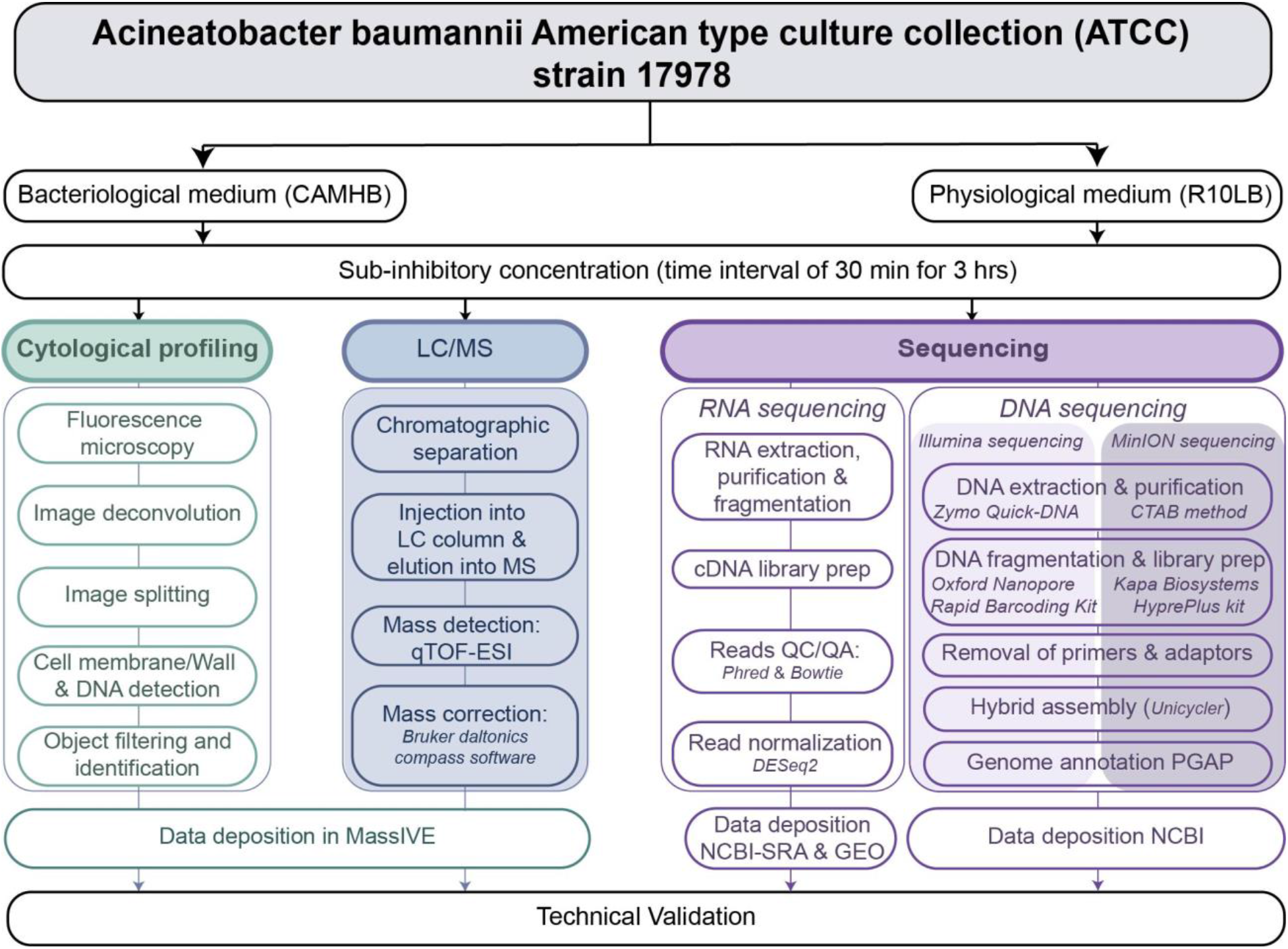
Diagram depicting the methodology used to profile the response of *Acinetobacter baumannii* strain ATCC 17978 to azithromycin in bacteriological and physiological media.

### DNA sequencing and genome assembly

Sequencing of the A. baumannii ATCC17978 reference genome was performed using an Illumina Hiseq 4000 (paired end, 150/150 bp reads) and ONT MinION to >50x coverage. The DNA for Illumina reads were prepared using a Zymo Research Quick-DNA Fungal/Bacterial Microprep Kit. Libraries were prepared using a Kapa Biosystems HyperPlus kit. The DNA for MinION sequencing was prepared by collecting high molecular weight genomic DNA using a CTAB method. Libraries were prepared using the ONT Rapid Barcoding Kit. Quality control steps were performed to remove unincorporated primers, adaptors, and detectable PCR primers prior to assembly. The complete genome was assembled into 3 contigs (genome and plasmids) using the “default” mode in Unicycler 0.4.2 and annotated using the NCBI Prokaryotic Genome Annotation Pipeline (PGAP) v4.11. The average Phred quality score for sequence reads is >32 which corresponds to a base calling accuracy of 99.99%. The reads are available at NCBI’s SRA under accession numbers SRR11671348 and SRR11671347. The complete genome is deposited at NCBI under accession numbers CP053098-CP053100. The final chromosomal genome size is 4,005,343 base pairs and the final plasmid sizes are 13,409 and 11,302 base pairs.

### cDNA library preparation and RNA sequencing

For RNA sequencing, 3 mL of culture was removed at mid-log phase (2.5 hrs of growth) and added to tubes which contained 6 mL of RNAprotect. Samples were incubated, centrifuged, and RNA was extracted using a ‘Quick RNA Fungal/Bacterial Microprep’ kit developed by Zymo Research. Cells were lysed with a Roche MagNa Lyser, DNase I was added, and RNA quality was determined using an Agilent Bioanalyzer. Ribosmal RNA was removed using an Illumina Ribo-Zero kit and then a cDNA library was created. A KAPA Stranded RNA-seq Library Preparation Kit was used for RNA fragmentation, sequencing adapter ligation, and library amplification. cDNA libraries were then sent for Illumina sequencing on a HiSeq 4000.

### RNA sequencing analysis

The reads were aligned to ATCC17978 strain genome using Bowtie2—. Alignment percentage was calculated by Bowtie2 while the average phred scores were calculated from bam files using samtools statistics method^20^. Read counts were next calculated using HTSeq counts package in ‘intersection-strict’ mode^21^. Using DESeq2, aligned reads were normalized to transcripts per million (TPM)^22^. The sklearn package was used to perform Principal Component Analysis (PCA) on log2(TPM + 1)^23^. The procedural steps for RNA sequencing analysis are presented in Fig. 2.

### Untargeted Liquid Chromatography Mass Spectrometry Data Acquisition

At the same time that samples of A. baumannii ATCC 17978 were taken for OD_600_ measurements, approximately 400 μL was collected from each replicate of all growth conditions and syringe-filtered using a 0.22 μm disc filters (Millex-GV, MilliporeSigma) to remove the cells from the spent media. Filtered samples were immediate placed on dry ice and then stored at −80 °C until liquid chromatography mass spectrometry (LC/MS) was performed. The LC/MS platform utilized an UltiMate3000 HPLC system (Thermo Scientific) paired to a ImpactHD (Bruker Daltonics) quadrupole-time-of-flight mass spectrometer. Filtered media from the RPMI+10% LB cultures were injected onto the LC at a volume of 5 μl, and filtered media from CA-MHB cultures were injected at a volume of 2 μl. All samples were injected onto a Kinetex 2.6μm polar-C18 reverse phase column (Phenomenex). The column temperature was maintained a 30 °C. For chromatographic separations, mobile phase A was LC/MS grade water modified with 0.1% formic acid and mobile phase B was LC/MS grade acetonitrile modified with 0.1% formic acid. Samples were injected at 95% A/5% B and at 1 minute the gradient was ramped to 65%A/35%B over the next 4 minutes. The solvent composition was stepped-up to 0%A/100%B and held for 1 minute before being restored to 95%A/5% for equilibration prior to injection of the subsequent sample.

Eluent from the HPLC was sprayed into the ImpactHD mass spectrometer via an Apollo II electrospray ionization source. The ImpactHD was controlled using otofControl v4.0.15 and the LC/MS sequence program was controlled using Hystar v3.2. During sample introduction into the mass spectrometer, the ESI source was configured to have a nebulizer gas pressure of 2 bar, drying gas flow rate of 9 liters/minute, and a drying gas temperature of 200°C. The mass spectrometer’s inlet capillary voltage was set at 3500 volts with an endplate offset of 500 volts. The mass spectrometer ion transfer optics were set to the following: Ion funnel 1 250 Vpp (volts peak-to-peak), ion funnel 2 250Vpp, transfer hexapole RF 100 Vpp, quadrupole ion energy 5 eV (electron volts), and collision quadrupole energy of 5 eV. The collision quadrupole RF was stepped across four voltages per scan: 450, 550, 800, 1100 Vpp. The collision cell transfer time was stepped across four values per scan: 70, 75, 90 and 95 μsecs. TOF pre-pulse storage was fixed at 7.0 μsecs. The mass spectrometer scan rate was fixed at 3Hz.

Prior to analysis, the mass spectrometer was externally calibrated using a sodium formate solution which was prepared by adding 100 μl of 1 M NaOH and 0.2% formic acid into 9.9mL of a 50%/50% water and isopropanol mixture. During mass spectrometric data acquisition, hexakis (1H,1H,2H-difluoroethoxy)-phosphazene (SynQuest Labs, Inc.) was used as a “lock mass” internal calibrant (m/z 622.028960; C_12_H_19_F_12_N_3_O_6_P_3_+). Subsequent to data acquisition, the lock mass was used to apply a linear mass correction to all mass spectra using Bruker Daltonics Compass Data Analysis software (ver. 4.3.110). Lock mass corrected data files were converted from the proprietary format (.d) to the mzXML open data format. All data herein were deposited to MassIVE (http://massive.ucsd.edu).

### Data Records

The reference genome assembly for *A. baumannii* ATCC 17978 was submitted in NCBI under accession number CP053098-CP053100 and the corresponding reads were submitted under accession number SRR11671348 and SRR11671347. The optical density and the corresponding calculated growth rates as well as the minimal inhibitory concentration measurements are available on Figshare. The BCP, HPLC, and Mass spectrometry data have been deposited to MassIVE repository under accession number MSV000085320. The complete RNAseq pipeline can be found on Figshare, Fastq files for each run have been deposited on NCBI-Sequence Read Archive. The overall summarized statistics for RNAseq and the metadata for all multi-omics data types are available on Figshare.

## Technical Validation

### Bacterial Cytological profiling

The image segmentations output by the CellProfiler pipeline was used to ensure accurate cell and object traces and measurements for representative images. “Parent” tags were used to match the cell outlines to related structures (e.g. DNA). The output files, which contain the identified cellular features, were uploaded to the MassIVE repository. Fig. 3 depicts a schematic of the image analysis pipeline for BCP data.

**Figure 3.**
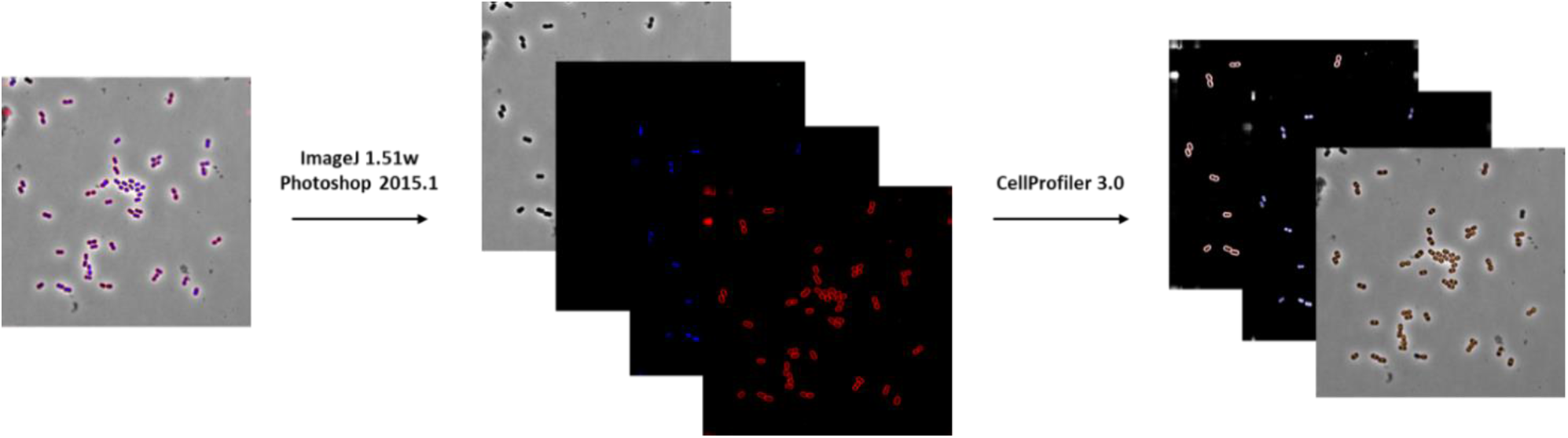
Bacterial Cytological Profiling (BCP) pipeline for *Acinetobacter baumannii* strain ATCC 17978.

### Untargeted Liquid Chromatography Mass Spectrometry data acquisition

For a given growth media type, each sample analyzed by LC/MS was largely comprised of the same molecules, a number of which were known. Therefore, extracted ion chromatograms (EICs) of these known molecules could be compared to evaluate the reproducibility of the retention time and ion intensity between experimental replicates. This reproducibility could then be evaluated by comparing both the overall base peak chromatogram (BPC) of experimental replicates, as well as EICs and peak areas of individual known molecules between experimental replicates. The EICs of known molecules were found to have a retention time drift of less than 0.1 minute across all samples and peak areas of known molecules differed by less than 15% between experimental replicates.

### RNA sequencing

An average of 94.41% of reads across all the samples aligned to the reference genome. The biological replicates showed high reproducibility with Spearman’s correlation coefficient >0.933 (Fig. 4).

**Figure 4.**
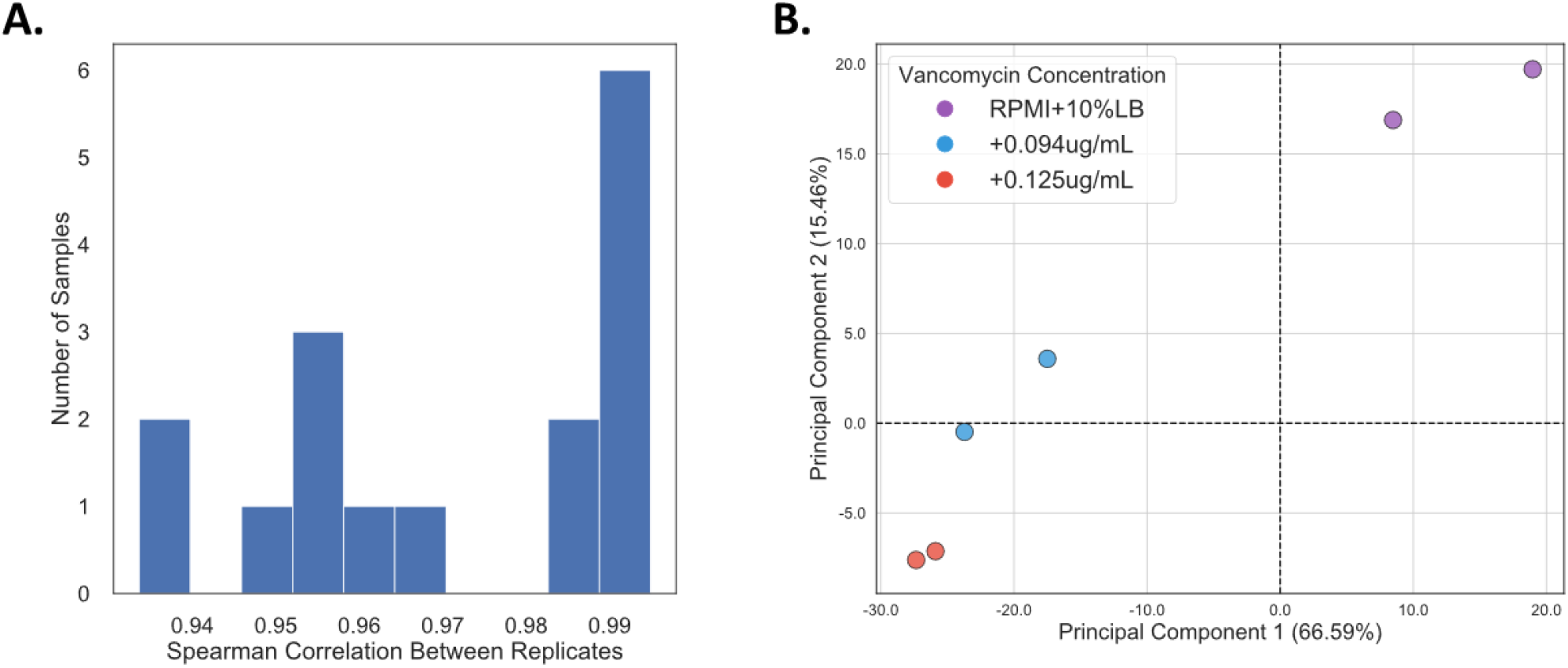
Quality control for transcriptomics sequencing data. (A) Histogram of Spearman’s correlation coefficient for TPM values between biological replicates. (B) PCA plot for expression profiles across all samples. Each sample point is color coded by azithromycin concentration to which the cultures were exposed and shaped according to the culture media type.

**Figure 5.**
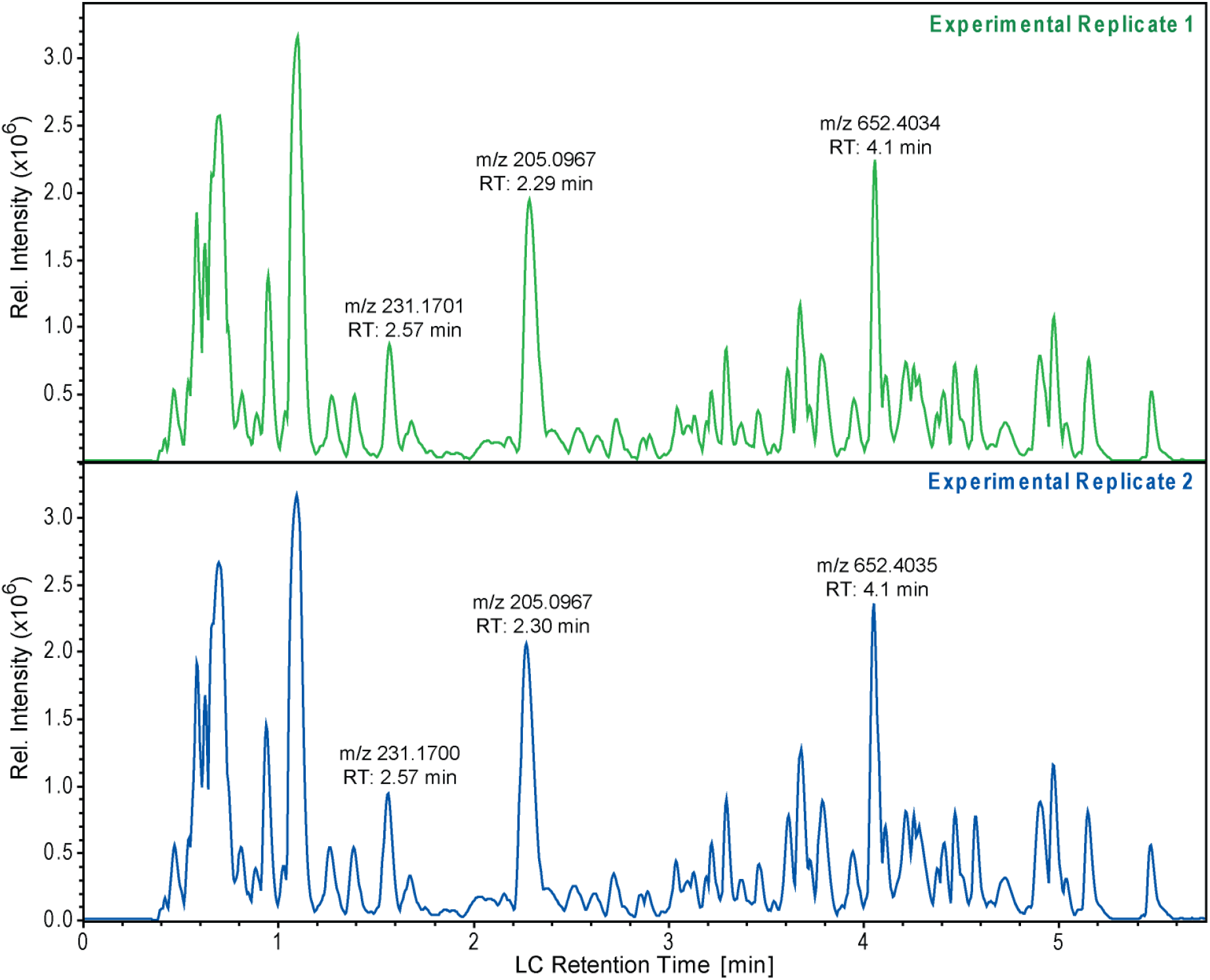
Validation of LC/MS base peak chromatograms of experimental replicates. Evaluation of the reproducibility of both retention time and peak areas within the LC/MS data were performed by direct comparison of the base peak chromatograms of experimental replicates. The plot represents a comparison of two separate experimental replicates of the same time-point from the same type of sample, in this case untreated ATCC 17978 grown in RPMI + 10% LB at 30 min.

## Code availability

The complete RNAseq pipeline used in analysis of RNAseq data is available on Figshare.

## Acknowledgements

We thank Anand Sastry for helping build the RNA sequencing analysis pipeline. This research was supported by NIH NIAID grant (1-U01-AI124316).

## Author Contributions

N.D. established the foundational growth and MIC experiments, compiled, analyzed results, and composed the manuscript. S.P. analyzed the RNA sequencing data and wrote Methods. H.T. performed growth experiments, analyzed the BCP data, and wrote Methods subsections. A.R. reviewed the article. M.M. analyzed the HPLC data and wrote subsections of the Methods. R.S. prepared samples for RNA sequencing and libraries for DNA sequencing and wrote Methods subsections. A.L. performed growth experiments and analyzed BCP data. A.V. performed growth experiments. J.S. wrote Methods. S.M.D. performed growth experiments. B.O.P and V.N. conceptualized the project, obtained funding, and edited the final manuscript.

## Bibliography

1 Antunes, L. C., Visca, P. & Towner, K. J. Acinetobacter baumannii: evolution of a global pathogen. Pathog Dis 71, 292–301, doi:10.1111/2049-632X.12125 (2014).

2 CDC. Antibiotic resistance threats in the United States. (Atlanta, GA: U.S. Department of Health and Human Services, 2019).

3 CDC. Antibiotic resistance threats in the United States. (US department of health and human services, 2013).

4 E. Tacconelli, N. & Magrini. Global priority list of antibiotic-resistant bacteria to guide research, discovery, and development of new antibiotics. WHO Press, 7 (2017).

5 Tacconelli, E. et al. Discovery, research, and development of new antibiotics: the WHO priority list of antibiotic-resistant bacteria and tuberculosis. The Lancet Infectious Diseases 18, 318–327, doi:10.1016/s1473-3099(17)30753-3 (2018).

6 Appleman, M. D. et al. In vitro activities of nontraditional antimicrobials against multiresistant Acinetobacter baumannii strains isolated in an intensive care unit outbreak. Antimicrob Agents Chemother 44, 1035–1040, doi:10.1128/aac.44.4.1035-1040.2000 (2000).

7 Nelson, R. E. et al. Costs and Mortality Associated With Multidrug-Resistant Healthcare-Associated Acinetobacter Infections. Infect Control Hosp Epidemiol 37, 1212–1218, doi:10.1017/ice.2016.145 (2016).

8 Leungtongkam, U., Thummeepak, R., Tasanapak, K. & Sitthisak, S. Acquisition and transfer of antibiotic resistance genes in association with conjugative plasmid or class 1 integrons of Acinetobacter baumannii. PLoS One 13, e0208468, doi:10.1371/journal.pone.0208468 (2018).

9 Spellberg, B., Bartlett, J., Wunderink, R. & Gilbert, D. N. Novel approaches are needed to develop tomorrow’s antibacterial therapies. Am J Respir Crit Care Med 191, 135–140, doi:10.1164/rccm.201410-1894OE (2015).

10 Lin, L. et al. Azithromycin Synergizes with Cationic Antimicrobial Peptides to Exert Bactericidal and Therapeutic Activity Against Highly Multidrug-Resistant Gram-Negative Bacterial Pathogens. EBioMedicine 2, 690–698, doi:10.1016/j.ebiom.2015.05.021 (2015).

11 Dillon, N. et al. Surprising synergy of dual translation inhibition vs. Acinetobacter baumannii and other multidrug-resistant bacterial pathogens. EBioMedicine, doi:10.1016/j.ebiom.2019.07.041 (2019).

12 Valentine, S. C. et al. Phenotypic and molecular characterization of Acinetobacter baumannii clinical isolates from nosocomial outbreaks in Los Angeles County, California. J Clin Microbiol 46, 2499–2507, doi:10.1128/JCM.00367-08 (2008).

13 Nonejuie, P., Burkart, M., Pogliano, K. & Pogliano, J. Bacterial cytological profiling rapidly identifies the cellular pathways targeted by antibacterial molecules. Proc Natl Acad Sci U S A 110, 16169–16174, doi:10.1073/pnas.1311066110 (2013).

14 Poudel, S. et al. Characterization of CA-MRSA TCH1516 exposed to nafcillin in bacteriological and physiological media. Sci Data 6, 43, doi:10.1038/s41597-019-0051-4 (2019).

15 Rajput, A. et al. Profiling the effect of nafcillin on HA-MRSA D712 using bacteriological and physiological media. Sci Data 6, 322, doi:10.1038/s41597-019-0331-z (2019).

16 Seif, Y. et al. Profiling the effect of nafcillin on HA-MRSA D592 using bacteriological and physiological media. bioRxiv, 2020.2004.2030.070904, doi:10.1101/2020.04.30.070904 (2020).

17 Poudel, S. et al. Revealing 29 sets of independently modulated genes in Staphylococcus aureus, their regulators and role in key physiological responses. bioRxiv, doi:10.1101/2020.03.18.997296 (2020).

18 Carpenter, A. E. et al. CellProfiler: image analysis software for identifying and quantifying cell phenotypes. Genome Biol 7, R100, doi:10.1186/gb-2006-7-10-r100 (2006).

19 Rodenacker, K. & Bengtsson, E. A feature set for cytometry on digitized microscopic images. Anal Cell Pathol 25, 1–36, doi:10.1155/2003/548678 (2003).

20 Anders, S., Pyl, P. T. & Huber, W. HTSeq--a Python framework to work with high-throughput sequencing data. Bioinformatics 31, 166–169, doi:10.1093/bioinformatics/btu638 (2015).

21 Li, H. et al. The Sequence Alignment/Map format and SAMtools. Bioinformatics 25, 2078–2079, doi:10.1093/bioinformatics/btp352 (2009).

22 Love, M. I., Huber, W. & Anders, S. Moderated estimation of fold change and dispersion for RNA-seq data with DESeq2. Genome Biol 15, 550, doi:10.1186/s13059-014-0550-8 (2014).

23 Johnson, W. E., Li, C. & Rabinovic, A. Adjusting batch effects in microarray expression data using empirical Bayes methods. Biostatistics 8, 118–127, doi:10.1093/biostatistics/kxj037 (2007).

